# Anti-glutamatergic effects of three lignan compounds: arctigenin, matairesinol and trachelogenin - An ex vivo study on rat brain slices

**DOI:** 10.1101/2022.10.14.512227

**Authors:** Peter Kiplangat Koech, Gergely Jócsák, Imre Boldizsár, Kinga Moldován, Sándor Borbély, Ildikó Világi, Arpád Dobolyi, Petra Varró

## Abstract

Arctigenin is a bioactive dibenzylbutyrolactone-type lignan exhibiting various pharmacological activities. The neuroprotective effects of arctigenin were demonstrated to be mediated via inhibition of AMPA/KA type glutamate receptors in the somatosensory cortex of the rat brain. The aim of this study was to compare the effects of arctigenin with matairesinol and trachelogenin on synaptic activity in *ex vivo* rat brain slices. Arctigenin, matairesinol and trachelogenin were isolated from *Arctium lappa, Centaurea scabiosa* and *Cirsium arvense*, respectively, and applied on brain slices via perfusion medium at the concentration range of 0.5-40 μM. The effects of the lignans were examined in the CA1 hippocampus and the somatosensory cortex by recording electrically evoked field potentials. Arctigenin and trachelogenin caused a significant dose-dependent decrease in the amplitude of hippocampal population spikes (POPS) and the slope of excitatory postsynaptic potentials (EPSPs), whereas matairesinol (1 μM and 10 μM) decreased EPSP slope but had no effect on POPS amplitude. Trachelogenin effect (0.5 μM, 10 μM, 20 μM) was comparable to arctigenin (1 μM, 20 μM, 40 μM) (p > 0.05). In the neocortex, arctigenin (10 μM, 20 μM) and trachelogenin (10 μM) significantly decreased the amplitude of evoked potential early component, while matairesinol (1 μM and 10 μM) had no significant effect (p>0.05). The results suggest that trachelogenin and arctigenin act via inhibition of AMPA/KA receptors in the brain and trachelogenin has a higher potency than arctigenin. Thus, trachelogenin and arctigenin could serve as lead compounds in the development of alternative neuroprotective drugs.

## Introduction

Lignans are a diverse group of phenolic compounds consisting of two phenylpropane units (C_6_-C_3_) linked by the side chain C-8, C-8’carbon atoms (Fig. 1) [1]. They are widely distributed in the plant kingdom and can be found in various plant organs, i.e., in fruits, flowers, seeds, roots, rhizomes, xylem, stems, leaves and resins. They are present in many plant foods, too. Recent research showed that consumption of dibenzylbutyrolactone lignans (DBLs) from plant foods may protect against cancer [2]. The closely related DBLs arctigenin (ATG), matairesinol (MAT) and trachelogenin (TGN) exhibit several biological activities, including neuroprotective, anticancer, antihypertensive, antiviral, antioxidant and anti-inflammatory effects [2]. As we confirmed recently, ATG, MAT and TGN can be isolated with high yield from the fruits of *Arctium lappa* L., *Asteraceae* [3,4], *Centaurea scabiosa* L., *Asteraceae* [3] and *Cirsium arvense* (L.) Scop, *Asteraceae* [5], respectively.

**Fig. 1.**
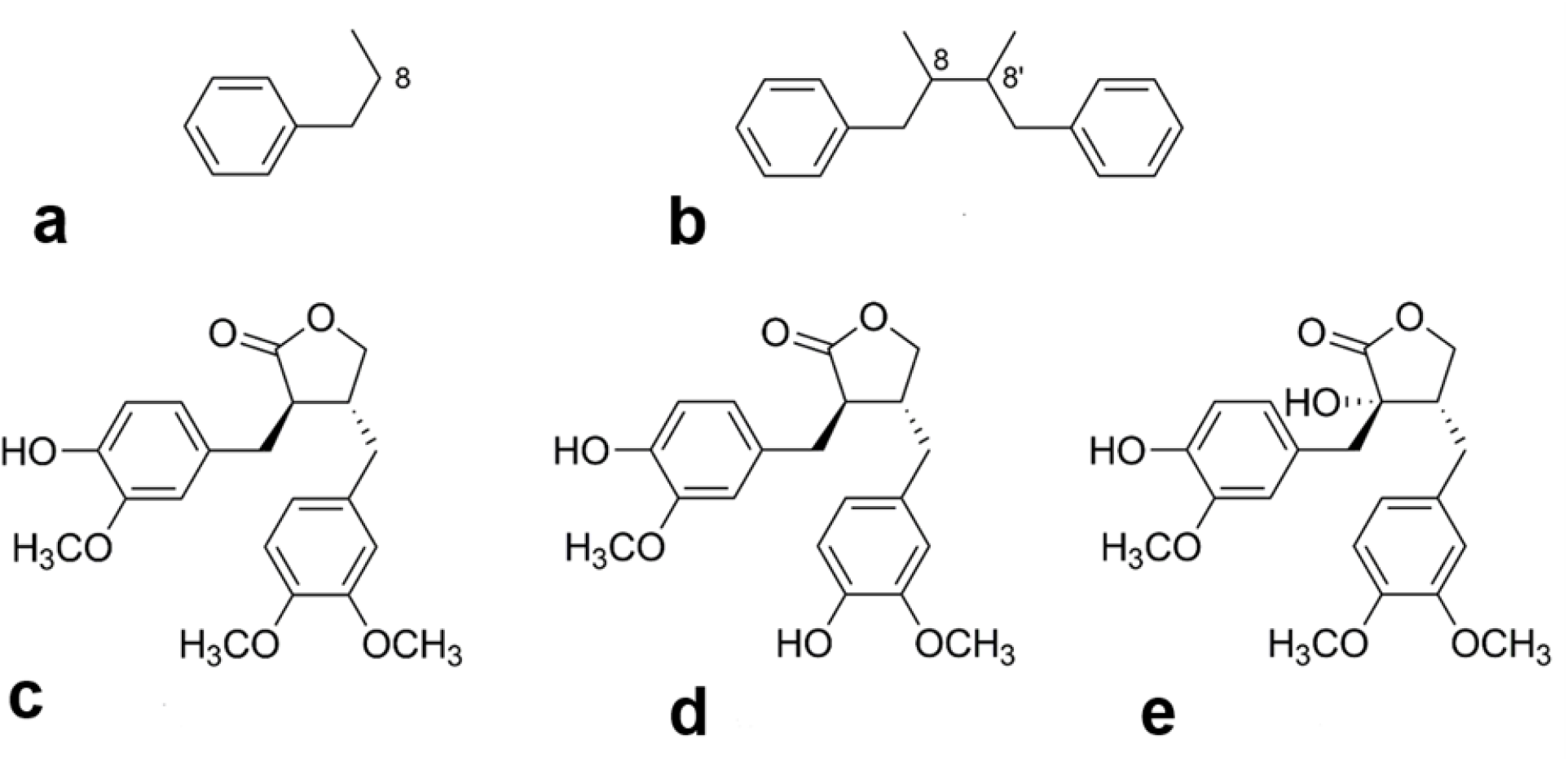
Chemical structure of lignans and the examined compounds. (**a**) The phenylpropane structural unit. **(b)** The basic chemical structure of lignans. (**c**) Structure of arctigenin (ATG). (**d**) Structure of matairesinol (MAT). (**e**) Structure of trachelogenin (TGN).

Glutamate is the major excitatory neurotransmitter in the brain that mediates synaptic transmission and plays a role in synaptic plasticity. Extracellular glutamate levels are low and tightly regulated. Thus, abnormally high extracellular glutamate levels lead to excitotoxicity by the overactivation of ionotropic glutamate receptors. This results in excessive Ca^2+^ influx into the cells, which activates a signaling cascade that ultimately leads to cell death [6].

Glutamate binds to AMPA (α-amino-3-hydroxy-5-methyl-4-isoxazolepropionic acid), kainate and NMDA (N-methyl-D-aspartate)-type receptors which are tetrameric ligand-gated ion channels. AMPA receptors are abundant and localized postsynaptically [7,8]. Kainate receptors are localized pre-, post-, and extra-synaptically, where they function in modulation of glutamate release, postsynaptic depolarization and regulation of intrinsic excitability [9]. They are abundant in the hippocampus and in layers 5-6 of the neocortex [10,11]. NMDA receptors are highly expressed in the hippocampus and cerebral cortex, located postsynaptically [12]. AMPA and kainate receptors primarily mediate Na^+^ influx, while NMDA receptors allow the passage of large amounts of Ca^2+^ [6]. AMPA receptors function as synaptic glutamate sensors and are responsible for rapid synaptic transmission [7,8]. NMDA receptors play a crucial role in synaptic plasticity, e.g. potentiation and memory processes [13]. Although kainate receptors play a minor role in mediating synaptic currents, they have been linked to fine-tuning synaptic efficacy and several brain disorders [14].

The high degree of plasticity of the hippocampus, especially that of the CA1 subregion, is associated with marked susceptibility to neurodegeneration, epilepsy, ischemia and chronic stress [15]. Neurodegenerative disorders are characterized by gradual loss of neurons resulting in deficits in memory, cognition and movement [16]. Epilepsy, Alzheimer’s and Parkinson’s disease are the most common brain disorders whose pathogenicity is linked to excitotoxicity [6,7]. Epilepsy is characterized by abnormal neuronal excitability and affects an estimated 50 million people worldwide, of which mesial temporal lobe epilepsy, associated with hippocampal sclerosis, is the most common type [17].

Neuroprotection by glutamate receptor antagonists is an essential pharmacological intervention for glutamate-induced excitotoxicity. In several disease models, blocking excessive AMPA receptor activation by inhibitors is neuroprotective [18]. However, early quinoxalinediones failed to cross the blood-brain barrier [7]. The applicability of AMPA receptor antagonists such as NBQX (2,3-dioxo-6-nitro-7-sulfamoyl-benzo[f]quinoxaline) is limited because they bind to all AMPA receptor subunits and kainate receptors [18]. In addition, NBQX precipitates in the kidneys and CNQX (6-cyano-7-nitroquinoxaline-2,3-dione) has limited ability to penetrate the blood-brain barrier [19]. 2,3-benzodiazepines are more selective for AMPA receptors but lack subunit selectivity [18]. 1-Naphthylacetylspermine, which inhibits only Ca^2+^-permeable AMPA receptors, also inhibits other ligand-gated ion channels [18].

Moreover, NMDA receptor antagonists block all the NMDA receptor activity leading to neurological adverse effects [20]. Hence, a search for alternatives, such as polyphenols with neuroprotective potential is ongoing.

The neuroprotective effect of arctigenin has been demonstrated in primary cultures of rat cortical cells and the somatosensory cortex [10,21]. ATG is lipophilic that can cross the blood-brain barrier and acts via AMPA/KA receptors [10,21]. TGN and MAT are molecularly related to ATG. TGN is a more potent Ca^2+^ antagonist than ATG, as demonstrated by inhibitory effects on K^+^-evoked contraction using isolated guinea pig taenia coli and subsequent determination of affinity values using spiral strips of rat aorta [22]. As we demonstrated recently, effects of TGN on rat ileal motility is about twice as potent as ATG [23]. Thus, we can assume that TGN and MAT will have more potent or similar effects on glutamatergic synapses in the brain as ATG.

This study aimed to determine and compare the anti-glutamatergic activities of ATG, TGN and MAT on *ex vivo* rat brain slices using electrophysiological methods.

## Results

Perfusion of the slices with pure ACSF or ACSF with 0.2% DMSO as vehicle for 30 min caused a slight increase in evoked potential amplitude in the hippocampus. ATG and TGN perfusion led to well observable, dose-dependent signal amplitude decreases, while MAT caused no significant change. At a higher concentration (40 μM), ATG caused a near-complete inhibition of evoked potentials, which was similar to that achieved by 10 μM CNQX (Fig. 3a).

**Fig. 2.**
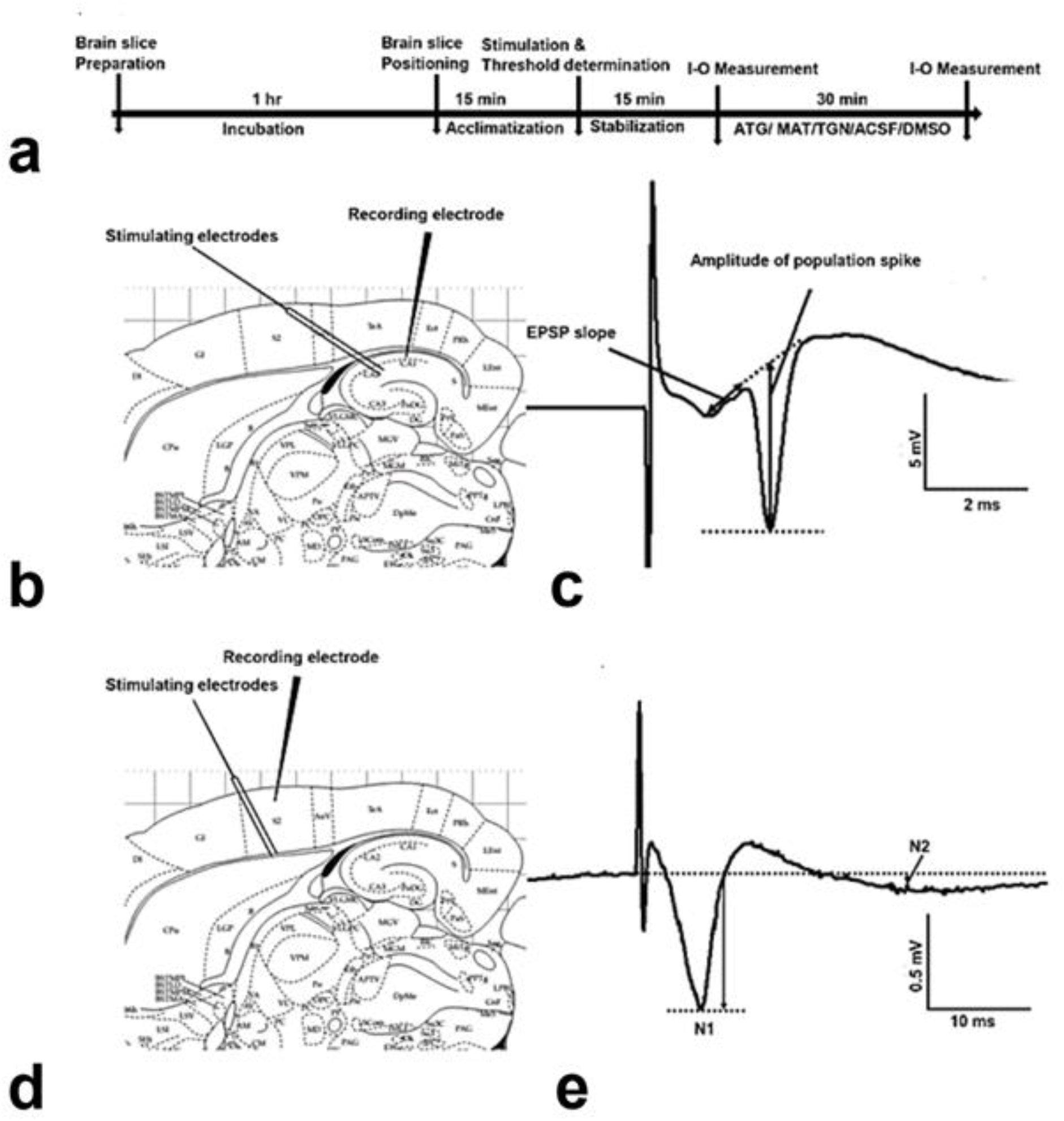
Design of the experiments in brain slices. **(a)** Time course of the experimental protocol. (**b**) Drawing of a living brain slice showing stimulating and recording electrode positions at Schaffer collaterals and *stratum pyramidale* of the CA1 hippocampus, respectively. **(c)** Original recorded signal in CA1 hippocampus characterized by EPSP slope and POPS amplitude. (**d)** Drawing of a living brain slice showing stimulating and recording electrode positions at greywhite matter border and layer 2/3 of the somatosensory cortex, respectively. **(e)** The original record of an evoked field potential in the neocortex characterized by the amplitudes of early (N1) and late (N2) negative peaks.

**Fig. 3.**
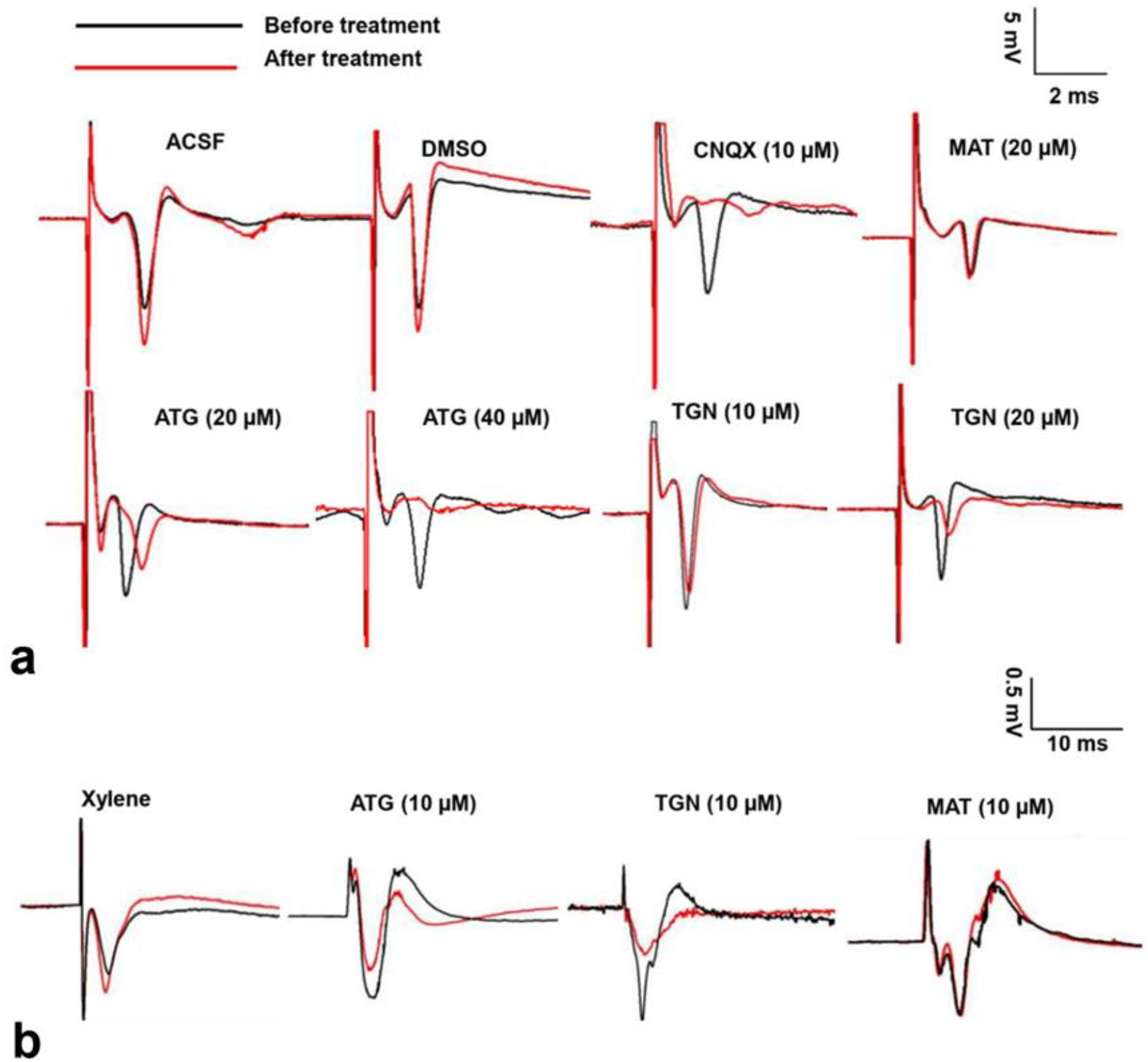
Effect of 30 min perfusion of brain slices with vehicle, different lignans or antagonist on evoked field potentials. (**a)** Representative evoked field potentials recorded at *stratum pyramidale* of CA1 hippocampus, showing the effects before (black line) and after (red line) treatment with ACSF, DMSO, 5 μM CNQX, 10 μM CNQX, 10 μM TGN, 20 μM TGN, 10 μM MAT, 20 μM MAT, 20 μM ATG and 40 μM ATG. (**b)** Representative evoked field potentials in the neocortex showing the effects before (black line) and after (red line) treatment with ACSF, 10 μM ATG, 10 μM TGN and 10 μM MAT.

In hippocampal slices, ATG and TGN caused a dose-dependent decrease in EPSP slope and POPS amplitude. 10 μM ATG, 20 μM ATG, 40 μM ATG, 10 μM TGN and 20 μM TGN lowered the EPSP slope and POPS amplitude significantly (p<0.01). 1 μM TGN significantly lowered the EPSP slope (p<0.01) but did not affect POPS amplitude (p>0.05). However, ACSF, DMSO, 1 μM ATG and 0.5 μM TGN had no effect (p>0.999). Interestingly, MAT showed significant inhibition of EPSP slope at lower concentrations of 1 μM and 10 μM (p<0.05) but did not significantly decrease the slope at a higher concentration of 20 μM (p>0.05). MAT did not significantly affect the POPS amplitude, either, at 1 μM, 10 μM and 20 μM (p>0.05). The effects of 40 μM ATG and 20 μM TGN on EPSP slope and POPS amplitude were comparable to that of 10 μM CNQX (p>0.999) (Fig. 4a, b).

**Fig. 4.**
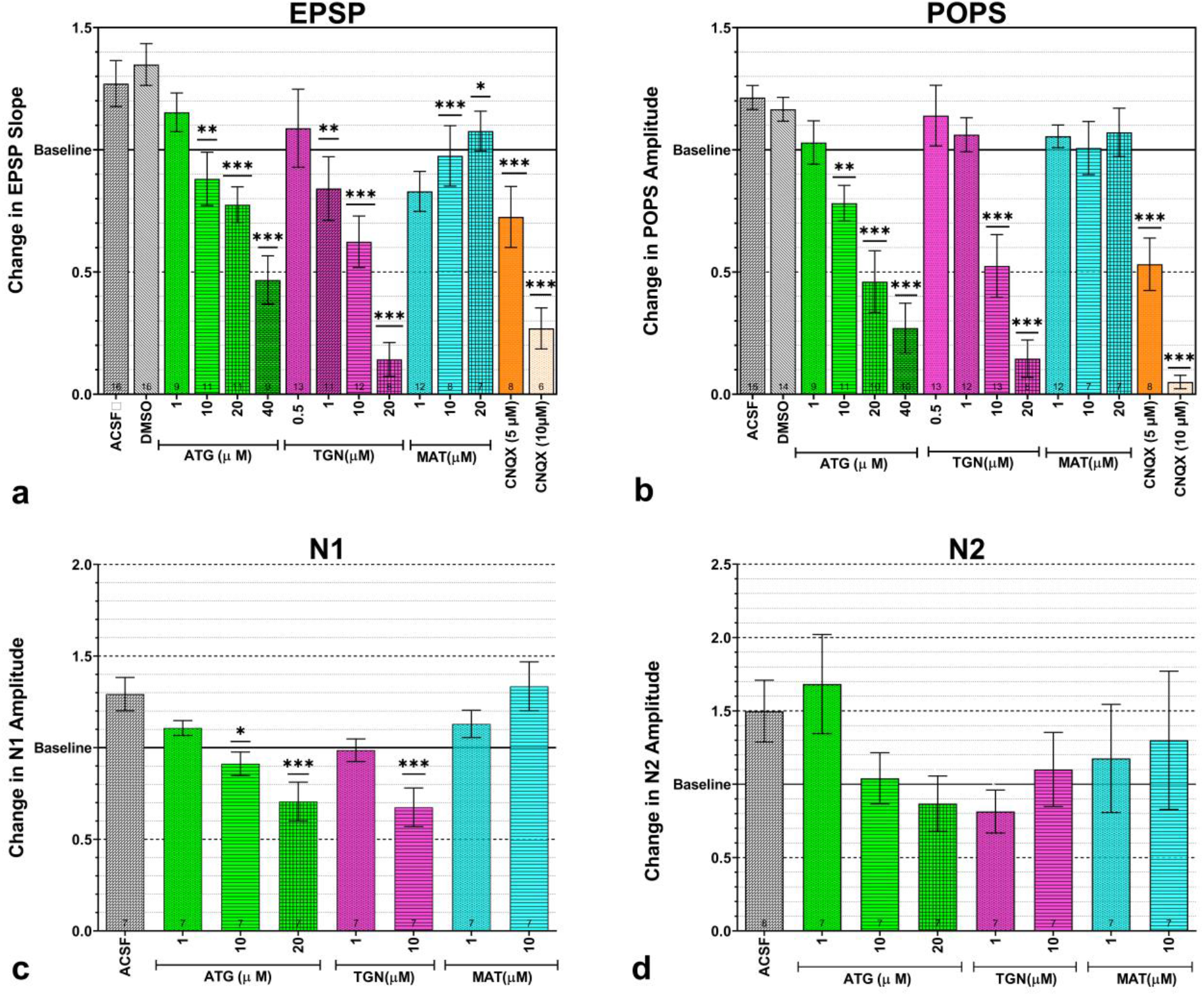
Effect of vehicle, different lignans or antagonist on the change in evoked field potential parameters. (**a**) ATG and TGN decreased EPSP slope in the CA1 hippocampus in a dose-dependent manner while MAT decreased the slope at 1 μM and 10 μM but not at 20 μM (**b**) ATG and TGN showed a dose-dependent decrease in POPS amplitude in the CA1 hippocampus while MAT had no effect. (**c**) In somatosensory neocortex, ATG and TGN decreased the N1 component amplitude dose-dependently while MAT had no effect. (**d**) ATG, TGN and MAT did not affect N2 amplitude. Bars represent means ± SEM of at least 7 independent measurements. *: p<0.05, **: p<0.01, and ***: p<0.001 indicate significant differences from DMSO control. Numbers within columns show sample numbers.

Regarding the neocortex, there were also observable changes in activity after treatment, as demonstrated by the evoked field potentials. ACSF slightly increased the baseline values of the early component, ATG and TGN decreased its amplitude, and MAT had no effect (Fig. 3b).

1 μM ATG and 1 μM TGN did not exert a significant inhibition on the increase in amplitude of N1 caused by ACSF (p>0.999). ATG and TGN caused dose-dependent decrease in N1 amplitude, which was significant in case of 10 and 20 μM ATG and 10 μM TGN (p<0.01). The degree of N1 amplitude suppression exerted by 10 μM TGN was comparable to that of 20 μM ATG (p>0.999). MAT (1 and 10 μM) did not affect N1 amplitude (p>0.999).

The N2 component was not significantly affected by any of the lignan treatments, but a tendency to inhibition can be observed similarly to the effect on N1 (Fig. 4c, d).

The effect of lignans on the shape of the input-output curves (I-O curves) was also analyzed. In case of hippocampal baseline recordings, the EPSP slope showed a linear increase from T to 3T, while POPS amplitude increases already saturated around 2.5T. The inhibitory effects exerted by lignans and CNQX manifested typically stronger at higher stimulation intensities, just like the slight excitatory effect of ACSF and DMSO perfusion. 20 μM TGN decreased the EPSP slope at 1.75T-3T stimulation intensities (p<0.05). Although the 20 μM ATG decreased the EPSP slope at 2.75T and above, the change was insignificant (p>0.05). 20 μM ATG, 20 μM TGN and 10 μM CNQX lowered the POPS amplitude at 1.25T to 3T (p<0.05). However, MAT (20 μM) had no effect on EPSP slope and POPS amplitude at all stimulation intensities (T-3T) (p>0.999) (Fig. 5A).

**Fig. 5.**
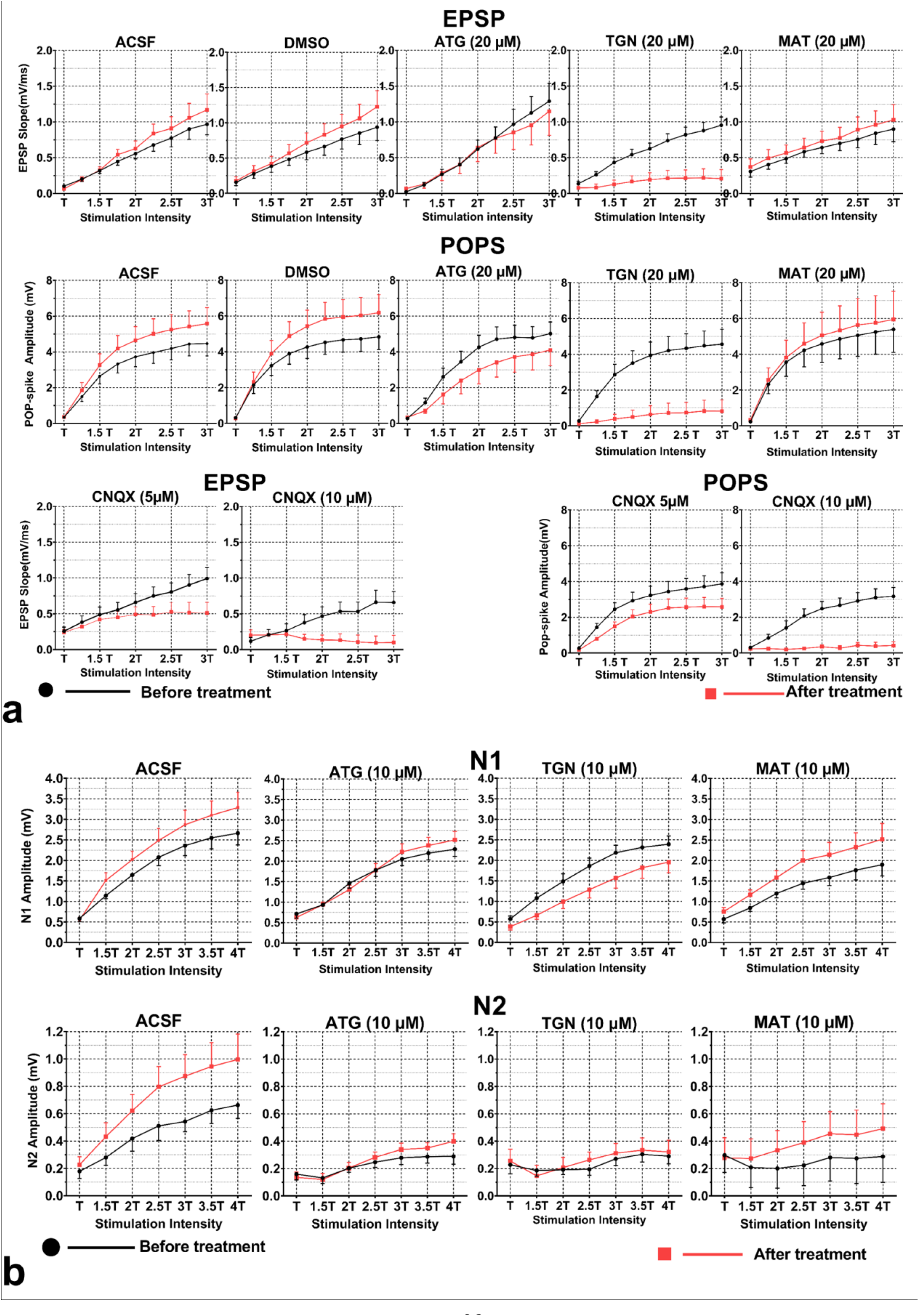
Effects of vehicle, different lignans or antagonist on input-output functions. (**a**) Input-output (I-O) curves in CA1 hippocampus showing the effects on EPSP slope and POP-spike amplitude before (black labels) and after (red labels) treatment with ACSF, ATG, TGN and MAT. Effects of DMSO and CNQX as negative and positive controls are also shown. (**b**) Input-output (I-O) curves in the neocortex showing the effects of ACSF, ATG, TGN and MAT on N1 and N2 amplitudes of the evoked response.

In neocortex, ACSF caused an increase in N1 and N2 amplitudes, which tended to grow with stimulus intensity, but it was not a significant change compared to the baseline (p>0.05). 20 μM ATG suppressed the N1 amplitude at 1.5T and 2T (p<0.01) but had no effect in other stimulation intensities (p>0.999). 10 μM TGN decreased the N1 amplitude at 1.5T to 4T (p<0.05). However, 1 μM ATG, 1 μM TGN, 1 μM MAT and 10 μM MAT had no effects at any stimulation intensity (p>0.999). The N2 component was not affected by any of the treatments, at any stimulation intensity (p>0.999) (Fig. 5b).

TGN is more potent than ATG as the effect of 0.5 μM TGN, 10 μM TGN and 20 μM TGN was comparable to 1 μM ATG, 20 μM ATG and 40 μM ATG, respectively (p>0.999), on both POP-spike amplitude and EPSP slope. Moreover, 20 μM TGN inhibited the EPSP slope to a higher extent than 20 μM ATG (p<0.001). The effect of 10 μM TGN on POP-spike amplitude and EPSP slope was comparable to 5 μM CNQX (p>0.999) (Fig. 4a,b). Similarly, in the neocortex, the inhibitory effect of 10 μM TGN on N1 amplitude was comparable to that of 20 μM ATG (p>0.999) (Fig. 4c). MAT was less potent than ATG and TGN and decreased EPSP slope at 1 μM and 10 μM but did not affect POPS, or N1 and N2 amplitudes in the neocortex. Doubling the concentration of MAT to 20 μM did not result in a significant change in EPSP slope and POPS amplitude (Fig. 4).

## Discussion

The effect of the dibenzylbutyrolactone lignan ATG was previously demonstrated to be mediated in rat somatosensory cortex via AMPA and Glu-K1 subunit-containing kainate receptors [10]. Its neuroprotective effects have been shown in primary cultures of rat cortical neurons [21]. A recent study reported a reversal of the abnormally high frequency of AMPA-mediated spontaneous excitatory postsynaptic currents to normal levels after ATG treatment [24]. Given the importance of AMPA and kainate receptors in the pathophysiology of many neurological disorders, we investigated whether ATG, TGN and MAT affect the electrically evoked field potentials of the rat CA1 hippocampus and neocortex *ex vivo*. Our previous hypothesis was that test lignans would block AMPA and kainate receptors, reducing postsynaptic depolarization and action potential initiation.

To determine the effects of lignans, evoked field potentials were tested before and after perfusion with lignan-containing ACSF and the change in activity relative to the baseline was determined. ATG and TGN caused a dose-dependent decrease in both EPSP slope and POP-spike amplitude in hippocampus and evoked field potential early component amplitude in neocortex. TGN already suppressed EPSP slope at a low concentration of 1 μM and the effect of 20 μM TGN was comparable to that of 40 μM ATG, indicating a higher potency for TGN compared to ATG. On the other hand, MAT did not have any significant effect in neocortex even at a high concentration of 20 μM, suggesting a weaker effect compared to ATG. Its suppressing activity in hippocampus at 1 μM on the EPSP slope and lack of effect on POPS amplitude indicates that the inhibition is insufficient to block action potentials. Surprisingly, this effect disappeared at higher concentrations.

To test the activity of lignans on brain slices, well-characterized brain areas were chosen in which glutamatergic pathways and field potentials have been already extensively analyzed. The field EPSP responses in hippocampus or neocortex indicate depolarization of the pyramidal neurons caused by synaptic activity and the steepness or amplitude of the EPSP slope is proportional to the amount of excitatory synaptic input. Measurement of the population spike amplitude in hippocampus estimates the number of neurons that reach firing threshold due to synaptic input [25]. In the neocortex, whereas mainly AMPA/kainate receptors mediate early synaptic evoked field response components, the late components are largely mediated by NMDA receptors [10].

Thus, the decrease in EPSP amplitude/slope mediated by the lignans indicates a decrease in glutamatergic synaptic input on the target cells, leading to reduced action potential firing. The effects of ATG and TGN were similar to that of the specific AMPA/kainate receptor antagonist CNQX, 20 μM TGN having about the same inhibitory effect as 10 μM CNQX. The lack of effect of all lignans on the late component of the neocortical evoked EPSP indicates the absence of a specific effect on NMDA receptors. Altogether, similar effects of the lignans could be demonstrated in both examined brain areas. The present results are consistent with previous studies in the somatosensory cortex, where 10 μM and 20 μM ATG reduced the evoked potential early component amplitude [10].

TGN and MAT are structurally closely related to ATG as they possess a five-membered lactone ring [26] and are also referred to as lignan-β-β’lactones [27]. TGN is structurally similar to ATG except for a hydroxyl group at C-8’ in TGN, replacing a hydrogen moiety in ATG. MAT differs from ATG at C-4, where the methoxy group is replaced by a hydroxyl group [28]. Previous studies showed that the number and arrangement of the phenolic hydroxyl group are crucial for the activity of lignans [29] and changing the position of a hydroxyl group leads to different biological responses [30]. The different position and number of the methoxy and hydroxyl groups could be the reason for the different activity of the test lignans.

Our results show that the antiglutamatergic activity of TGN was the highest among the three examined compounds, being about twice stronger than that of ATG. The high activity is presumably due to a hydroxyl group on the butyrolactone ring, which is absent in both ATG and MAT [28]. In a recent study, the glutamate receptor antagonistic effects of ATG and its analogues were studied *in vitro* [31]. According to their results, removing methoxy or hydroxyl groups from the phenolic rings (rendering it less polar) increases the compound’s activity. Similar observations were made *in vivo*, investigating insect feeding deterrent activity: non-polar substituents like the methoxy group appear to increase the lignans’ biological activity, while polar substituents like the hydroxyl group decrease it [32]. Thus, the lower potency of MAT compared to ATG in our study could be due to the presence of a hydroxyl group instead of a methoxy group.

Finally, TGN and ATG had similar effects as CNQX, suggesting that anti-glutamatergic effects are mediated by AMPA/kainate but not NMDA receptors. The extent of inhibition is concentration-dependent, decreasing the synaptic input and postsynaptic depolarization. This leads to a decreased synaptic strength and an inability to generate action potentials, as shown by electrophysiological results.

## Conclusion

Plants possess numerous secondary metabolites, which are important sources of new drugs. Plants belonging to the Asteraceae family are rich sources of lignans. Although dietary lignans are potentially beneficial to health, *in vivo* effects of the test lignans present in various foods have not been sufficiently investigated. Lignans exhibit a wide range of biological activities, including neuroprotective effects. We hereby report that dibenzylbutyrolactone lignans inhibit synaptic activity via AMPA/kainate receptors. TGN is more potent than ATG, but MAT is less potent than ATG and TGN. Thus, ATG and TGN represent potential lead compounds for developing alternative neuroprotective drugs that could potentially be used in various brain disorders, such as epilepsy, schizophrenia, depression, bipolar disorder, and mental retardation.

## Materials and Methods

### Drugs and Chemicals

Artificial cerebrospinal fluid (ACSF) with the following composition (mM): 126 NaCl; 26 NaHCO_3_; 1.8 KCl; 1.25 KH_2_PO_4_; 1.3 MgSO_4_; 2.4 CaCl_2_; 10 glucose (pH=7.4) was prepared at the Department of Physiology and Neurobiology, Eötvös Loránd University. Cyanquixaline (6-cyano-7-nitroquinoxaline-2,3-dione; CNQX; >99%) was purchased from Hello Bio Ltd. Dimethyl sulfoxide (DMSO; ≥99.9%) was used for dissolution of lignans. Unless stated otherwise, all drugs were purchased from Sigma-Aldrich Ltd.

### Isolation and formulation of the arctigenin, matairesinol and trachelogenin

Fruits of *Arctium lappa, Centaurea scabiosa* and *Cirsium arvense, Asteraceae* were collected from different Hungarian habitats. Their voucher specimens were deposited in the herbarium of the Department of Plant Anatomy, Eötvös Loránd University, Budapest, Hungary. Plant materials have previously been identified by Dr. Imre Boldizsár (Department of Plant Anatomy, Eötvös Loránd University) [3,4,5]. Plant names have been checked with http://www.theplantlist.org. ATG, MAT and TGN are accumulated in the fruits of *A. lappa, C. scabiosa* and *C. arvense* in their glycoside form connected to a glucose molecule [3,4,5]. The fruits of *A. lappa* and *C. arvense* also contain endogenous glycosidase enzymes allowing the enzymatic hydrolysis of ATG and TGN glycosides in water medium at 40 °C, resulting in the formation of free aglycones ATG and TGN [4,5,33]. However, glycosidase enzymes responsible for the hydrolysis of MAT glycoside are not present in the fruits of *C. scabiosa* [33]. Consequently, to isolate the free aglycones (ATG, MAT, TGN), enzymatic hydrolysis of ATG and TGN glycosides in the fruits of *A. lappa* and *C. arvense*, and acidic hydrolysis of MAT glycoside in the fruit of *C. scabiosa*, need to be performed.

### Endogenous enzymatic hydrolysis of ATG and TGN glycosides in the fruits of A. lappa and C. arvense

Lyophilized and pulverized *A. lappa* and *C. arvense* fruit samples (2.0 g) were suspended in 5.0 mL of distilled water. These suspensions were heated at 40°C for 60 min., allowing for enzymatic conversions of ATG and TGN glycosides in an aqueous medium. Thereafter, the samples were lyophilized and extracted.

### Extraction

Lyophilized and pulverized intact fruit sample of *C. scabiosa* (2.0 g), as well as enzyme-treated and lyophilized tissues of *A. lappa* and *C. arvense* were extracted with 25 mL of 80% (v/v) methanol held under reflux at boiling point for 60 min. Thereafter, the insoluble, centrifuged material was extracted for a second time, as before. The combined supernatants were dried by a rotary vacuum evaporator at 30–40°C.

### Acidic hydrolysis of MAT glycoside in C. scabiosa fruit extract

Dried extract prepared from the intact fruits of *C. scabiosa* was dissolved in 5.0 mL of 2 M trifluoroacetic acid (in 25 mL screw-capped vial). This solution was heated for 60 min at 100 °C. After heating, the sample was dried using a vacuum evaporator (at 30–40 °C). *Isolation by preparative high-performance liquid chromatography (HPLC*)

The dried extracts of the enzyme-hydrolyzed fruit samples (*A. lappa* and *C. arvense*) and the dried sample of the acid-hydrolyzed extract (*C. scabiosa*) were dissolved in 2.5 mL of 80% (v/v) methanol for the isolation of ATG, TGN and MAT by preparative HPLC as described in our previous manuscript [3]. Their structures were confirmed by HPLC hyphenated with ultraviolet (UV) and high-resolution mass spectrometry (HR-MS) detection (Fig. 1). Purity of isolated ATG and TGN was recently determined to be 95.4 % and 96.7 %, respectively [23] and that of MAT was determined to be 95.2% by a HPLC-UV-HRMS method (Supporting information, Fig. 1S).

The isolated compounds were dissolved in dimethyl sulfoxide (DMSO) to obtain 10 mM stock solution, then stored at −20°C. The stock solution was diluted in ACSF to obtain the final concentrations of ATG (1 μM, 10 μM, 20 μM and 40 μM), MAT (1 μM,10 μM, and 20 μM) and TGN (0.5 μM, 1 μM, 10 μM and 20 μM).

### Experimental animals

Adult male Wistar rats weighing 200-335 g were purchased from Toxi-coop Ltd. Before the experiments, they were kept under controlled laboratory conditions, 12:12-h light:dark photocycle and controlled temperature (22 ± 2°C). They were fed with standard rat pellets and tap water was provided *ad libitum*. Experiments were carried out per the Hungarian Act of Animal Care and Experimentation (1998, XXVIII) and with the directive 2010/63/EU of the European Parliament and of the Council of 22 September 2010 on the protection of animals used for scientific purposes. All procedures were reviewed and approved by the Animal Care and Use Committee of Eötvös Loránd University. All efforts were made to reduce the number of animals used and minimize animal suffering.

### Preparation of rat brain slices and electrophysiological recordings

The experimental rats were deeply anesthetized with chloral-hydrate (350 mg/kg i.p.) and decapitated with a rodent guillotine. The whole brain was quickly removed from the skull, the cerebellum and the olfactory bulb were excised, and the left and right cerebral hemispheres were separated lengthwise and mounted. Then, 400 μm thick horizontal slices containing the cortex and hippocampus were cut with a vibratome (OTS-4500, Electron Microscopy Sciences /Leica VT1000s, Leica Biosystem) under ice-cold ACSF, bubbled with carbogen (95% O2, 5% CO2). Before recording, the slices were incubated at room temperature for 1 hour in the same solution. Individual slices were then transferred to a Haas-type interface recording chamber (Experimetria Ltd) through which standard ACSF bubbled with carbogen was perfused via a peristaltic pump at a rate of 2.5 ml/min and maintained at 33°C ± 1°C. The brain slices for hippocampal experiments were acclimatized for 15 min in the recording chamber (Fig. 2a). During this time, tungsten bipolar stimulating electrodes (FHC) were positioned at the Schaffer collaterals and glass recording microelectrodes (5-10 MΩ) filled with 1M NaCl were positioned in the pyramidal cell layer (*stratum pyramidale*) of the CA1 region (Fig. 2b). Field potentials were evoked by electrical stimulation with 100 μs duration square voltage pulses, and bandpass-filtered signals (0.16Hz-1kHz) were amplified 1000-fold with an Axoclamp2A amplifier (Axon Instruments) and digitized with an A/D converter (NI-6023E A/D card, National Instruments). Before recording, the viability of the slice was tested by examining if the amplitude of evoked population spikes (POPS) reached 1.5 mV. The evoked field potentials consisted of the POPS sitting on a field EPSP (excitatory postsynaptic potential), and analyzed parameters were POPS amplitude and the initial slope of the EPSP (Fig. 2C). The stimulation voltage threshold (T) was determined, and the slices were stabilized at 2T for 15 min, stimulated every 10 s, and T was adjusted again after the stabilization period. An input-output (I-O) curve was recorded by gradually increasing the stimulus intensity in 8 steps from T to 3T in 0.25T increments, with stimulation every 10 s. Then, 10 evoked responses were recorded at 2T. The brain slices were then perfused with ACSF containing different concentrations of arctigenin, trachelogenin or matairesinol, DMSO (vehicle control), CNQX (positive control), or pure ACSF (negative control) for 30 min stimulated at 2T, every 20 s. An I-O curve and 10 evoked responses at 2T, stimulated every 10 s, were recorded again (after treatment condition) (Fig. 2a). In the case of the neocortical slices, field potentials were evoked by positioning the recording electrode in layer 2/3 of the somatosensory cortex and stimulating electrodes just below the recording electrode at the white-grey matter border (Fig. 2d). The viability of brain slices was tested before recording. If the peak-to-peak amplitude of the maximal evoked response was less than 1 mV, the slice was excluded from the experiment. The recording protocol was the same as for hippocampal slices, the stimulation intensities applied for the I-O curve recoding being slightly different: here, the stimulus intensity was gradually increased from T to 4T in 6 steps in 0.5T increments. Analyzed parameters of the evoked field responses were the amplitudes of early and late negative peaks (N1 and N2 amplitudes) (Fig. 2e). Here, the treatment groups were 1 μM or 10 μM MAT, TGN or ATG. All hippocampal and neocortical signals were recorded and analyzed using the Solution Pack for Experimental Laboratories (SPEL) Advanced Intrasys computer program (Experimetria Ltd).

### Statistical analysis

189 horizontal slices originating from 80 animals and at least 7 slices per treatment/control group were evaluated. The stored cortical and hippocampal signals were analyzed using SPEL advanced Intrasys computer program. Data were exported to OriginPro 8 to plot evoked field potential traces. Data were transformed and normalized using MS Excel for Windows and expressed as the mean and standard error of the mean (SEM). Data were verified for normal distribution with the Kolmogorov-Smirnov test. Differences between experimental/control groups were estimated by two-way ANOVA followed by Bonferroni *post hoc* test. All analyses and graphical representations were performed using Graph Pad Prism software version 9.1.2 for Windows. A p-value of less than 0.05 was considered statistically significant.

## Abbreviations

AMPA: α-amino-3-hydroxy-5-methyl-4-isoxazolepropionic acid
ACSF: artificial cerebrospinal fluid
ATG: arctigenin
CNQX: 6-cyano-7-nitroquinoxaline-2,3-dione
DBL: dibenzylbutyrolactone lignan
DMSO: dimethyl sulfoxide
EPSP: excitatory postsynaptic potential
HPLC: high performance liquid chromatography
I-O: input-output
MAT: matairesinol
NBQX: 2,3-dioxo-6-nitro-7-sulfamoyl-benzo[f]quinoxaline
NMDA: N-methyl-D-aspartate
POPS: population spike
SEM: standard error of the mean
T: voltage stimulation threshold
TGN: trachelogenin

## Supporting Information

HPLC-UV chromatogram of matairesinol and methodical details for its isolation is available as Supporting Information.

## Acknowledgments

This work was supported by the Hungarian National Research, Development, and Innovation Office (grant numbers OTKA K-135712, OTKA K-134221 and National Brain Research Program grant NKFIH-4300-1/2017-NKP_17). This work was completed in the framework of the ELTE Thematic Excellence Programme 2020 supported by the Hungarian National Research, Development, and Innovation Office (TKP2020-IKA-05). We are thankful to the Stipendium Hungaricum Scholarship Programme for financial support to Peter Kiplang’at Koech. Sponsors had no involvement in study design, in the collection, analysis and interpretation of data, in the writing of the report and in the decision to submit the article for publication. We are grateful for the technical assistance of Gábor Tolmár.

## Conflict of interest

The authors declare no conflict of interest.

## Notes

### Competing Interest Statement

The authors have declared no competing interest.

