## Supplemental material for "Anti-glutamatergic effects of three lignan compounds: arctigenin, matairesinol and trachelogenin - An ex vivo study on rat brain slices"

***Dr. Petra Varró***

Department of Physiology and Neurobiology, Institute of Biology

Eötvös Loránd University

Pázmány Péter sétány 1/C

1117 Budapest

Hungary

Supplementary Fig. 1S shows the HPLC-UV chromatogram of the isolated matairesinol, confirming its excellent purity (95.2%).

Instruments, eluents and detection parameters of the HPLC separation were identical to those reported in our recent article (Berek-Nagy et al., Phytochemistry 190 (2021) 112851). A linear gradient program (0.0 min, 20% B; 15.0 min, 70% B; 19.0 min 90% B) was used. Column: Kinetex C18 column (75 × 3 mm; 2.6 μm) (Phenomenex, USA). Flow rate: 0.4 mL/min, injected volume: 2 µL.

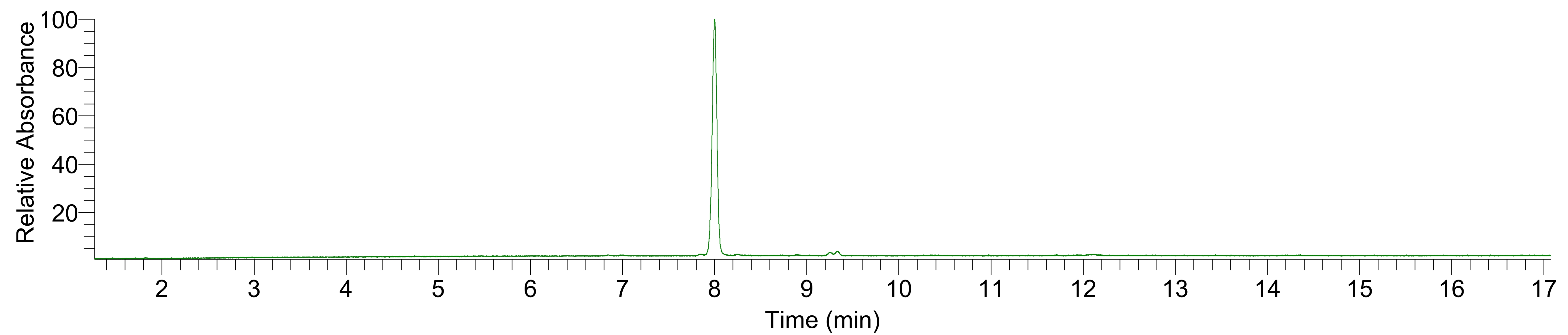

**Fig. 1S.**

HPLC-UV (λ=280 nm) chromatogram of matairesinol dissolved in methanol (concentration: 0.05 mg/mL).
